# Strength of low-frequency EEG phase entrainment to external stimuli is associated with fluctuations in the brain’s internal state

**DOI:** 10.1101/2024.05.23.595436

**Authors:** Verónica Mäki-Marttunen, Alexandra Velinov, Sander Nieuwenhuis

## Abstract

The brain attends to environmental rhythms by aligning the phase of internal oscillations. However, the factors underlying fluctuations in the strength of this phase entrainment remain largely unknown. In the present study we examined whether the strength of low-frequency EEG phase entrainment to rhythmic stimulus sequences varied with pupil size and posterior alpha-band power, thought to reflect arousal level and excitability of posterior cortical brain areas, respectively. We recorded pupil size and scalp EEG while participants carried out an intermodal selective attention task, in which they were instructed to attend to a rhythmic sequence of visual or auditory stimuli and ignore the other perceptual modality. As expected, intertrial phase coherence (ITC), a measure of entrainment strength, was larger for the task-relevant than for the task-irrelevant modality. Across the experiment, pupil size and posterior alpha power were strongly linked with each other. Interestingly, ITC tracked both variables: larger pupil size was associated with a selective increase in entrainment to the task-relevant stimulus sequence, whereas larger posterior alpha power was associated with a *decrease* in phase entrainment to both the task-relevant and task-irrelevant stimulus sequences. Exploratory analyses showed that a temporal relation between ITC and posterior alpha power emerged in the time periods around pupil maxima and pupil minima. These results indicate that endogenous sources contribute distinctly to the fluctuations of EEG phase entrainment.

**Significance statement:** Fluctuations in cortical state powerfully shape the perception of external stimuli. Understanding the physiological signatures of cortical state fluctuations is crucial to understand how the brain selectively attends and switches between internal and external content. Here we studied how two signatures of attentional state, pupil-linked arousal and power in the alpha band, shape the entrainment of brain activity to low-frequency rhythmic stimuli. Our results reveal common and dissociable influences of these signatures at slow time scales. Furthermore, measuring and including pupil size and posterior alpha power as covariates in statistical models can help increase statistical power in studies focusing on EEG phase entrainment. Our study provides new evidence on a direct influence of cortical state on the perception of rhythmic stimuli.

## Introduction

How we perceive the continuous stream of external sensory stimulation is highly modulated by the internal state of the brain (Matthewson et al., 2012), but the interaction between external and internal mechanisms of attention is still poorly understood. Rhythmic stimuli entrain endogenous oscillatory activity in the brain to the temporal structure of the stimulus sequence (Bauer et al., 2018; Ten Oever et al., 2017; Van Rullen et al., (2014); Zoefel et al., 2018). This *neural entrainment* results in the alignment of the high-excitability oscillatory phase with the predicted onset of the rhythmic stimuli (Lakatos et al., 2008; Stefanics et al., 2010), and provides an advantage for the processing of natural stimuli that are inherently rhythmical, such as speech, music and repetitive motion (Henry and Obleser, 2012; Lakatos et al. 2019). Although phase alignment of oscillations to external rhythms constitutes an important principle of neural dynamics and attention, little is known about the factors that make the strength of phase entrainment fluctuate over time.

Here, we investigated the impact on low-frequency phase entrainment of two physiological measures known to be highly variable across time: pupil-linked arousal and posterior alpha-band power. Attention during tasks requiring sustained performance commonly fluctuates due to variations in the internal state of vigilance or arousal (Unsworth and Robinson, 2017). Pupil size has long been used to measure variations in arousal level in humans (Bradshaw, 1967). Very low or very high levels of vigilance, as characterized by lapses of attention or task-unrelated thoughts, often occur during periods of small and/or large (pre-trial) baseline pupil diameter (Madore et al., 2020; Unsworth and Robinson, 2016; van den Brink et al., 2016). Whether and how arousal modulates the strength of neural entrainment, which is an externally directed process, is still unknown.

Changes in posterior alpha-band power, as measured with electroencephalography (EEG), are thought to reflect the excitability of posterior cortical brain areas that process external (sensory) input (Romei et al., 2008; Samaha et al., 2016). Increased alpha power is suggested to facilitate internally directed attention rather than attention to external stimuli (Benedek et al., 2014; Cooper et al., 2003; Klimesch et al., 2007). Lapses of attention and task-unrelated thoughts are associated with increases in pre-trial posterior alpha power (Compton et al., 2019; Groot et al., 2021; Jin et al., 2019; Madore et al., 2020; O’Connell et al., 2009). Similarly, a study in nonhuman primates found that periods with high alpha power were characterized by attentional lapses and strongly diminished low-frequency entrainment of neuronal oscillations to task-relevant rhythmic stimulus sequences (Lakatos et al., 2016). These results suggest that alpha power is an index of internal state and would be expected to anti-correlate with the strength of entrainment to a task-relevant sequence of stimuli.

Although baseline pupil size and posterior alpha have both been associated with attentional state, their interrelation may not be straightforward. Alpha power as a measure of inattention, as reviewed above, would be expected to correlate negatively with pupil size. That is, low arousal would be related to distraction and an increase in the “idling” alpha. However, studies on the relationship between pupil size and posterior alpha power at rest or in pre-trial periods have found a positive relation and/or an inverted-U relationship (Ceh et al., 2020; Hong et al., 2014; Pfeffer et al., 2022; Podvalny et al., 2021; van Kempen et al., 2019, but see Waschke et al., 2019). Therefore, these two signals may reflect brain states that are at least partly dissociable, and we expected that they would distinctly influence neural entrainment.

In this study, we aimed to test whether and how low-frequency phase entrainment is susceptible to modulations in internal state, as indexed by pupil-linked arousal and posterior alpha power. We simultaneously recorded pupil size and scalp EEG to investigate the relationships between pupil-linked arousal, cortical alpha oscillations and low-frequency phase entrainment during an intermodal attention task. In the task, rhythmic streams of visual and auditory stimuli were concurrently presented. Participants had to detect infrequent targets in the cued modality and ignore the other modality. A small difference in repetition rate (visual, 1.1 Hz, auditory, 1.4 Hz) allowed us to separately assess entrainment to the task-relevant and task-irrelevant stimulus sequences. We hypothesized that transient changes in the strength of entrainment to the attended modality would be related to fluctuations in the arousal level of the participants. We also expected that internal neural processes signaled by alpha power would modulate entrainment level. This study brings novel insights into the interrelation between external and internal attentional mechanisms of the brain.

## Methods

### Participants

Thirty-seven students recruited at Leiden University took part in the experimental study. One participant was excluded because the recording was interrupted. EEG data from four participants were excluded from analysis due to excessive artifacts (> 50% of epochs rejected) in the Attend Visual condition. One participant was excluded because of low accuracy (hit rate < 0.5) in the Attend Auditory condition. Five out of the first 15 participants were excluded because they reported an unforeseen visual illusion, where they perceived the afterimage of the green fixation stimulus having a similar color as the magenta targets, probably caused by a complementary afterimage effect (Manzotti, 2017). As a precaution to avoid this illusion, after the first 15 participants we changed the color stimulus, as reported below. Performance did not differ between either set of stimuli for the participants included in the final study (hit rate, first round: 0.77 ± 0.05, second round: 0.79 ± 0.07, T = 0.86, p = 0.398; incorrect responses, first round: 32 ± 30, second round: 38 ± 38, T = 0.41, p = 0.685). The final sample consisted of 26 participants (mean age: 23, age range: 18-28; 22 female). Study exclusion criteria included any psychiatric disorders and wearing contact lenses. The experimental protocol was reviewed and approved by the ethics committee of the Authors’ University. Participants signed a consent form prior to participating and received a compensation of 7.5 euros/hour or course credits.

### Task and stimuli

Participants performed an intermodal selective attention task (**Fig. 1**), based on that used by Lakatos et al. (2016) to study phase entrainment in monkeys. Participants were presented with simultaneous streams of visual and auditory stimuli and had to respond as quickly as possible to rare target stimuli in the cued modality by pressing the “B” key on a keyboard, and to otherwise withhold responding. The response deadline was 1 second after target onset. The task consisted of ten blocks, each lasting approximately five minutes. The participants were prompted at the beginning of each block to attend to the visual stimuli (Attend Visual condition) or to the auditory stimuli (Attend Auditory condition), and ignore the other modality. Half of the blocks required the participant to attend to the visual modality, the other half to the auditory modality, in a quasi-random order (A1-V1-A2-A3-V2-A4-V3-V4-A5-V5). In each block 605 stimuli were presented: 265 visual stimuli and 340 auditory stimuli. In each stream of stimuli, 15% of the stimuli were targets and 85% were standards. The auditory stimuli consisted of short beeps with a 460-Hz pitch for the standards and a 476-Hz pitch for the targets. The visual stimuli consisted of colored RGB Gabor patches (orange standards: RGB 229 133 126, magenta targets: RGB 229 90 172, green fixation point for participants #1-15: RGB 151 199 62; purple fixation point for participants #16-37: RGB 180 137 205) with a 10-degree orientation and a frequency of 0.1 cycles/pixel, created using an online Gabor-patch generator (https://www.cogsci.nl/gabor-generator). Because changes in luminosity can drive changes in pupil size, we made sure that the visual stimuli and the inter-stimuli screens were isoluminant. For this, visual stimuli were made isoluminant with the grey screen background (RGB 167 167 167) using the SHINEcolor toolbox (Dal Ben, 2023). The visual and auditory stimuli were presented at average frequencies of 1.1 Hz and 1.4 Hz, respectively. All stimuli were presented for 50 ms. The fixation point was on the screen in the interval between consecutive visual stimuli, and all visual stimuli subtended a 4° angle on the screen. The average interstimulus interval was 387 ms, with a range from 0 (i.e., auditory and visual stimuli presented simultaneously) to 727 ms. To lower a possible contribution of stimulus-evoked EEG responses to our measure of intertrial phase coherence, we added to the interstimulus interval some decorrelated jitter, drawn on each stimulus time from the values [-66, -33, 0, 33, 66 ms]. Entrainment occurs at the average (i.e., expected) frequency of presentation of the stimuli, and thus is independent of the stimulus presentation itself (Besle et al., 2011).

**Figure 1.**
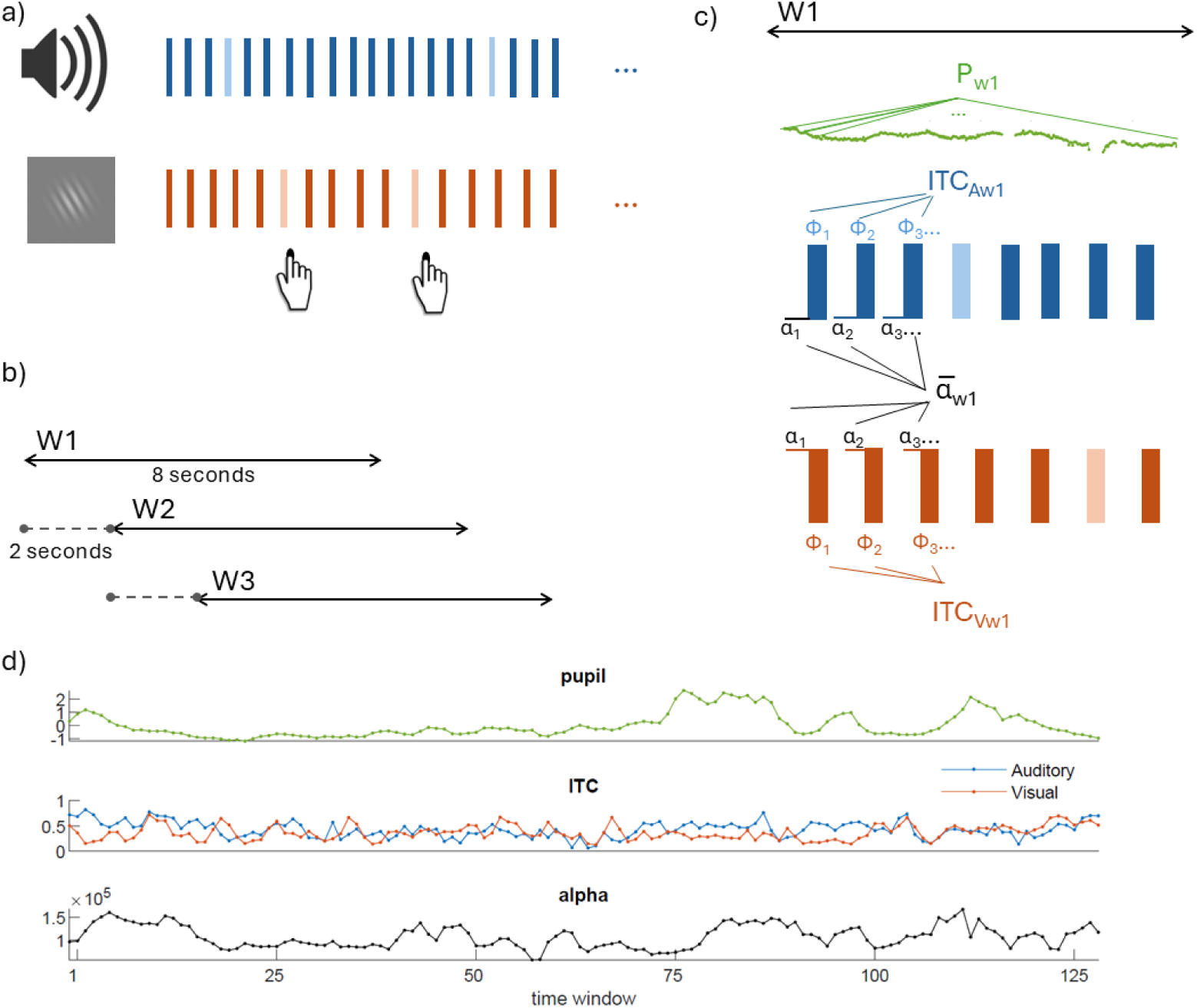
Task and analysis overview. a) Multimodal task. Toy example of a sequence of stimuli sequences of the intermodal selective attention task. Rectangles represent stimuli of the auditory modality (tones, represented in blue) and the visual modality (Gabor patches, represented in orange). Auditory and visual stimuli were presented simultaneously at slightly different frequencies (auditory: 1.4 Hz, visual: 1.1 Hz). Participants had to attend to only one modality per block, and respond to the target stimuli in that modality (represented in lighter color) with a button press. In this example, the participant responds to the target stimulus in the visual modalityꟷthat is, the visual stimuli were task-relevant. b) Illustration of sliding windows approach. Pupil size, ITC and alpha power were calculated in 8-seconds sliding windows (W1, W2, W3, etc) covering the whole block, with 2-seconds steps. c) Illustration of the calculation of ITC over auditory (ITC_A_) and visual (ITC_V_) stimuli types in one 8-seconds sliding window. Average pupil size (P, pupil trace in green) was calculated for the entire window. Alpha power for each window was calculated as the average pre-stimulus alpha power across all stimuli. ITC represented the coherence of phase values at a given frequency across all the trials of the same modality. d) A participant’s single-block sample data of time courses of the various measures, where each observation represents one time window. These time courses were used to study the relation between pupil, pre-stimulus alpha power, and ITC.

### Procedure

Participants were instructed to sit as still as possible throughout the blocks and to always look at the center of the screen, also in the Attend Auditory condition. Their chins were positioned in a chin rest, with their eyes at approximately 73 cm from a monitor with a refresh rate of 60 Hz. For the auditory stimuli custom in-ear air-tube headphones were used to prevent electrical artifacts in the EEG recordings. Participants first performed two practice blocks, one for each condition, which each lasted 2 minutes and contained a total of 256 visual and auditory stimuli. After the practice blocks, the eye-tracker was calibrated and the participants performed the task in a dimly lit room (19.9 lx). The task was controlled using Eprime 3.0 with the appropriate Tobii extensions installed. In between blocks, participants could take breaks and decide when to start the following block. Lighting conditions and screen luminance were kept constant across the whole session. Participants took between 60 and 70 minutes to complete the experiment.

### Time-windows approach

The purpose of this study was to examine the slow temporal relation between pupil-linked arousal, entrainment and alpha power. Based on a previous study (Lakatos et al. 2016), we analyzed the pupil and EEG data in sliding time windows. The windows had a length of 8 seconds and a step size of 2 seconds, resulting in an overlap of 6 seconds between consecutive time windows (**Fig. 1**). The window length and step size were chosen to achieve a balance between the number of stimuli included in each estimate of entrainment (larger number provides a better estimate) and the temporal resolution of our analyses (shorter window length provides higher resolution). Each time window included approximately nine visual stimuli and approximately eleven auditory stimuli.

Given that the paradigm involved the simultaneous presentation of visual and auditory stimuli, each time window contained stimuli in both modalities, but depending on the block, only one of them was task-relevant. For each time window we obtained the following five measures (**Fig. 1**):

(i) Average pupil size across the entire 8-second window.

(ii and iii) intertrial phase coherence (ITC, a measure of entrainment), which was calculated based on the phase angles at stimulus onset of all (ii) visual stimuli and all (iii) auditory stimuli of which the onset latency fell within the 8-second window.

(iv and v) Average pre-stimulus alpha power across all (iv) visual stimuli and all (v) auditory stimuli of which the onset latency fell within the 8-second window.

Note that in our key analyses we recoded the visual and auditory stimulus sequences as ‘task-relevant’ and ‘task-irrelevant’, based on the participant’s instruction, so that for each 8-second window we had task-relevant and task-irrelevant ITC and pre-stimulus alpha power values.

Below, we explain how pupil size, ITC and pre-stimulus alpha power were computed.

### Pupillometry

The pupil diameter of both eyes was recorded using a Tobii Pro X3-120 eye-tracker with a sampling rate of 40 Hz. The calibration was performed at the beginning of the task using a 5-point fixation procedure. The Tobii device automatically marked periods when no pupil signal was recorded (e.g., blinks) and custom code was compiled to save accurate timing data that could then be used to create eye-tracking epochs and segmentations for the behavioral data. Raw pupil data were preprocessed using the PhysioData Toolbox v0.5.0 (Kret and Sjak-Shie, 2019) using the standard pipeline of the Pupil Diameter Analyzer module. One participant was excluded from the pupil analyses due to excessive data loss during preprocessing (> 50% of recording). An exponential decay in pupil size was observed at the beginning of the blocks for most participants. Since the magnitude of such time-on-task effects can obscure the effects of the more subtle trial-by-trial fluctuations in pupil-linked arousal that we were interested in (van den Brink et al., 2016), we fitted an exponential decay curve to the pupil time series using the MATLAB R2018 (MathWorks, Inc., Natick, MA) functions *fittype* and *fit*, and regressed out this component from the pupil data. For the time windows analyses, the continuous pupil signal was cut into 8-second windows (2-second step size) and then averaged across each window. The resulting time series were *z*-scored separately for each block to make sure that differences in pupil size reflected local fluctuations in pupil size instead of global time-on task effects (Hopstaken et al., 2015).

### EEG acquisition and preprocessing

EEG data were recorded using a BioSemi ActiveTwo system with 32 active Ag-AgCl electrodes, at a sampling rate of 512 Hz. The electrodes were arranged according to the international 10-20 system. In addition to the 32 main electrodes, two reference electrodes were positioned on the mastoids, two facial electrodes above and below the left eye and two electrodes on the temples. Raw EEG data were preprocessed in EEGLAB (Delorme and Makeig 2004). The signal was re-referenced to the mastoids and high-pass filtered (0.5 Hz). Bad channels were removed using *cleanrawdata* function of EEGLAB. The minimal acceptable correlation with nearby channels was set to the default value of 0.8, while the parameter for artifact subspace reconstruction (a method for removing high-amplitude artifacts from EEG data) was set to 10. Missing channels were interpolated from neighboring sites. Then the data were split into Attend Auditory and Attend Visual conditions and epoched into 3-second segments centered on stimulus onset. This epoch length was chosen to avoid edge artifacts in the time-frequency analysis (Besle et al. 2011). Bad epochs were rejected automatically (threshold criteria: 150 uV) and an independent component analysis (infomax algorithm with *extended* option to extract sub-Gaussian sources) was performed to identify and remove noise components manually (e.g., eye artifacts, muscle artifacts). Preprocessing was done using the Neuroscience Gateway (https://www.nsgportal.org, Martínez-Cancino et al., 2021; Sivagnanam et al., 2013).

### Intertrial phase coherence and prestimulus alpha power

The epoched EEG data were used to calculate measures of phase entrainment and alpha power. We first extracted spectral power and phase dynamics using Morlet wavelet decomposition from 1 to 30 Hz with 30 logarithmically spaced steps. We used a wavelet cycle range of 3 to 12. This corresponded to a precision, as measured by the spectral full-width at half-maximum (FWHM), ranging from 0.48 to 5.88 Hz (Cohen, 2019). Entrainment of EEG oscillations to the stimulus rhythms is calculated based on the instantaneous phase values at the frequency and time of presentation of visual and auditory stimuli (Lakatos et al. 2013, Besle et al. 2011). To determine the phase angle at stimulus onset we used the 3-second epochs centered on stimulus onset that were mentioned above (section “EEG acquisition and preprocessing”). Our ITC estimates based on the instantaneous phase at t = 0 were not affected by edge-effect artifacts (i.e., were within the cone of influence). Intertrial phase coherence (ITC), a measure of the consistency across trials of the phase angle at the time of stimulus onset (Tallon-Baudry et al., 1996), was calculated with the following formula:

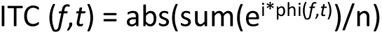

where phi is the phase at frequency *f* and time point *t* of a single trial (i.e. stimulus presentation), and n is the number of trials considered. This was done separately for the visual and auditory stimulus streams at the corresponding frequency (1.1 Hz for Visual and 1.4 Hz for Auditory, **Fig. 3a**). For these analyses, we averaged data from electrodes displaying the largest ITC for each modality: AF3, AF4, F3, Fz, F4, FC1 and FC2 for auditory stimuli and C3, Cz, C4, CP1 and CP2 for visual stimuli. For the 8-seconds time window analyses, we computed the ITC across all visual and (separately) auditory stimuli of which the onset latency fell within the 8-second window.

For the analysis of prestimulus alpha power, we extracted for each epoch the data from -500 to 0 msec before the stimulus and applied a fast Fourier transform to extract the power density spectrum (Compton et al. 2019; Ergenoglu et al. 2004). We then calculated the area under the curve for the alpha band (8-13 Hz). **Fig. 3-1** shows the posterior distribution of prestimulus alpha power. We averaged the data of posterior electrodes (P7, P3, Pz, O1, Oz, O2, P4 and P8), obtaining thus one prestimulus alpha value for each stimulus. For the 8-seconds time window analyses, we computed the average alpha power value across all visual and auditory stimuli of which the onset time fell within the 8-second window. The resulting time series of ITC and alpha power were *z*-scored separately for each block.

The analyses described above eventually yielded one pupil-size value for each 8-second window, and two ITC values and two alpha power values for each 8-second window: one for the task-relevant stimulus sequence (visual stimuli in the Attend Visual condition; auditory stimuli in the Attend Auditory condition) and one for the task-irrelevant stimulus sequence (visual stimuli in the Attend Auditory condition; auditory stimuli in the Attend Visual condition).

### Temporal analysis around pupil maxima and minima

We analyzed the variables ITC and prestimulus alpha power in segments of 10 windows around maxima and minima of the pupil timecourse. Because of the overlap between windows, 10 windows corresponded to 26 seconds. The normalized pupil size timecourse of each participant was thresholded at +1 standard deviation to detect the peaks and -1 standard deviation to detect the troughs. For this, the function *findpeaks* in MATLAB was employed, with a minimum distance between consecutive maxima and consecutive minima of 10 time windows. Then, the normalized ITC and alpha power data were extracted from those windows for further analysis. The cross-correlation between the variables was calculated with the function *xcorr* in MATLAB. To statistically compare cross-correlations between the time-windowed variables around pupil maxima and minima, we used cluster-based permutation testing, with 1000 iterations and a significance level of α = 0.01 for the individual time points and α = 0.05 for the significant clusters. All the temporal analyses were performed on the task-relevant modality, and data were collapsed across modalities.

### Statistics

Hit rate was defined as the proportion of targets in the cued modality that were detected in time (i.e., button press within 1 second after target onset). Any response that was not a hit was counted as an incorrect response. Given the overlap between the task-relevant and task-irrelevant stimulus sequences, it was hard to distinguish incorrect responses to standards in the task-relevant modality from responses to the targets of the task-irrelevant modality, and thus we pooled them together as ‘incorrect responses’. Average ITC values were compared with repeated-measures ANOVA with cued modality (Attend Visual, Attend Auditory) and stimulus type (visual, auditory) as within-subject factors. To examine the relationships between pupil size, ITC, prestimulus alpha power and task performance (hit rate, number of incorrect responses), we aggregated the data of the 8-second time windows in 10 equally populated bins on the basis of one variable, and examined the effects of bin (1-10), task relevance (task-relevant vs. task-irrelevant stimulus sequence) and stimulus type (visual, auditory) on one of the other variables using a repeated-measures ANOVA. In the analyses of task performance we included in the model both a linear and quadratic effect of bin. Statistics were performed using MATLAB.

## Results

### Behavioral results

Participants showed better performance in the Attend Auditory condition than in the Attend Visual condition (**Fig. 2a**). The hit rate was significantly higher (*F*(1, 495) = 6.78, *p* = 0.009), whereas the number of incorrect responses did not differ between conditions (*F*(1, 475) = -0.88, *p* = 0.11; for statistics, see Table 1). We also inspected the relation between pupil size and task performance. Pupil bin had no linear and quadratic effects on hit rate (*F*(1, 495) = 1.79, *p* = 0.18; *F*(1, 495) = 2.52, *p* = 0.11). In contrast, pupil bin had both a linear and a quadratic effect on the number of incorrect responses (*F*(1, 475) = 22.65, *p* < 0.001; *F*(1, 475) = 30.90, *p* < 0.001). More incorrect responses were made at intermediate levels of pupil size (**Fig. 2a**). No significant interactions between pupil size bin and cued modality were observed (hits: *p* = 0.64, incorrect responses: *p* = 0.74).

**Figure 2.**
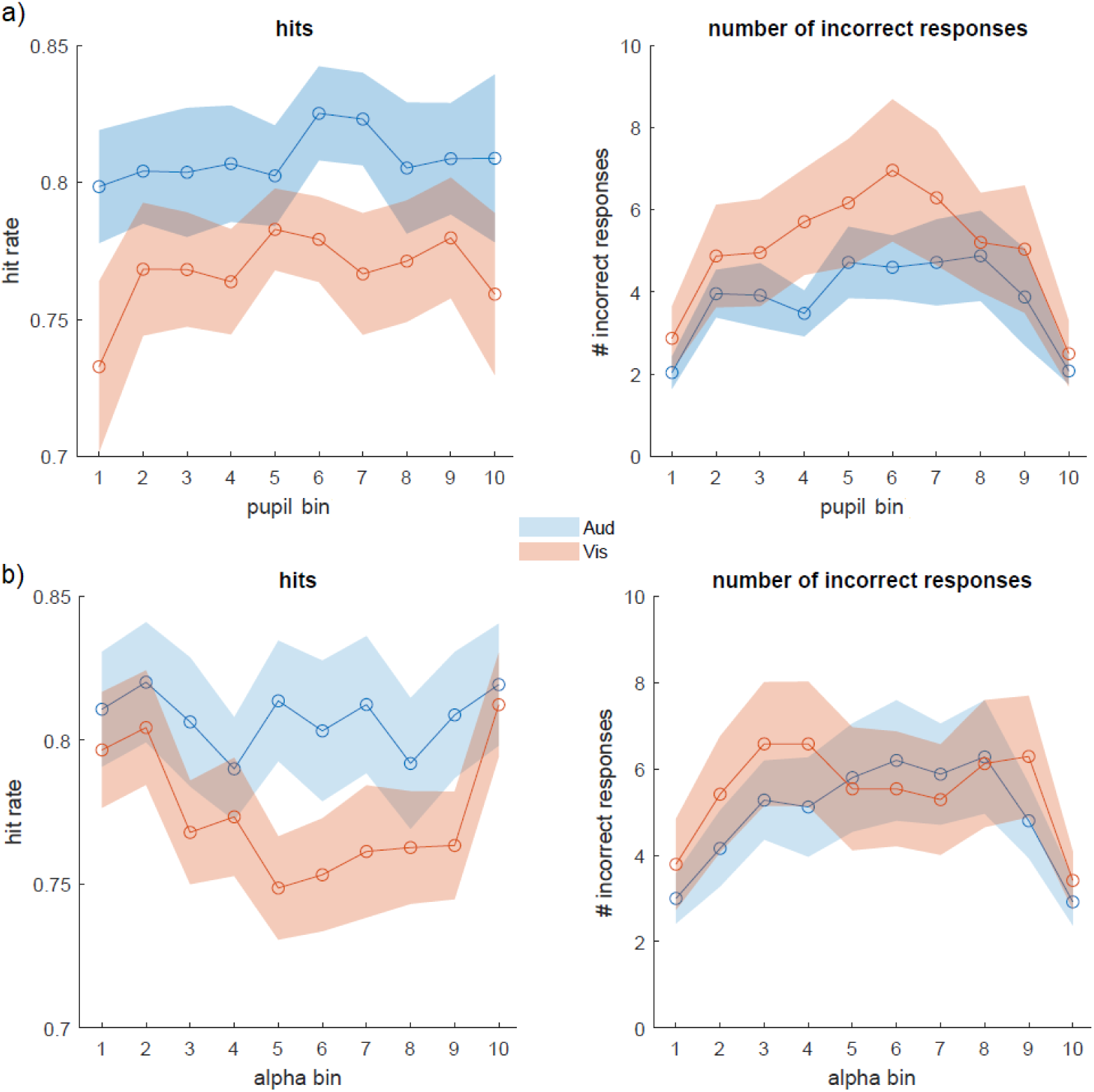
Behavioral results. Hit rate and number of incorrect responses as a function of pupil bin (a) and prestimulus alpha bin (b). Colours correspond to the different task-relevant modalities (Aud: auditory; Vis: visual).

**Table 1.** Statistical table.

Prestimulus alpha power showed a qualitatively similar relationship with task performance. Alpha bin had no linear and quadratic effects on hit rate (*F*(1, 515) = 0.04, *p* = 0.62; *F*(1, 515) = 0.08, *p* = 0.76). In contrast, alpha bin had both a linear and a quadratic effect on the number of incorrect responses (*F*(1, 495) = 15.678, *p* < 0.001; *F*(1, 495) = 19.135, *p* < 0.001). More incorrect responses were made at intermediate levels of prestimulus alpha power (**Fig. 2b**). No significant interaction between alpha power bin and cued modality was observed (hits: *p* = 0.16, incorrect responses: *p* = 0.89).

### Low-frequency phase entrainment to simultaneous trains of auditory and visual stimuli

**Fig. 3a** shows the ITC spectra for the various conditions. The top row shows strong ITC at the frequency corresponding to the task-relevant stimulus sequence (1.4 Hz for Attend Auditory, 1.1 Hz for Attend Visual), suggesting entrainment of EEG oscillations to the task-relevant stimulus rhythm. The corresponding ITC topography was frontal for the auditory stimulus sequence and central for the visual stimulus sequence (**Fig. 3b**). The bottom row of **Fig. 3a** suggests that EEG oscillations also became entrained to the task-irrelevant stimulus sequences, although significantly less than to the task-relevant stimulus sequences (**Fig. 3c**) and with more distributed topographies (**Fig. 3b**). The ITC associated with the task-relevant stimulus sequence was larger in the visual than in the auditory modality (F(1, 100) = 28.67, p < 0.001; Fig. 3c), and the difference between entrainment to task-relevant and task-irrelevant stimulus sequences was also larger in the visual modality (interaction task relevance * stimulus type: F(1, 100) = 7.68, p = 0.006; task-relevant vs. task-irrelevant: visual: p < 0.001, auditory: p = 0.056).

**Figure 3.**
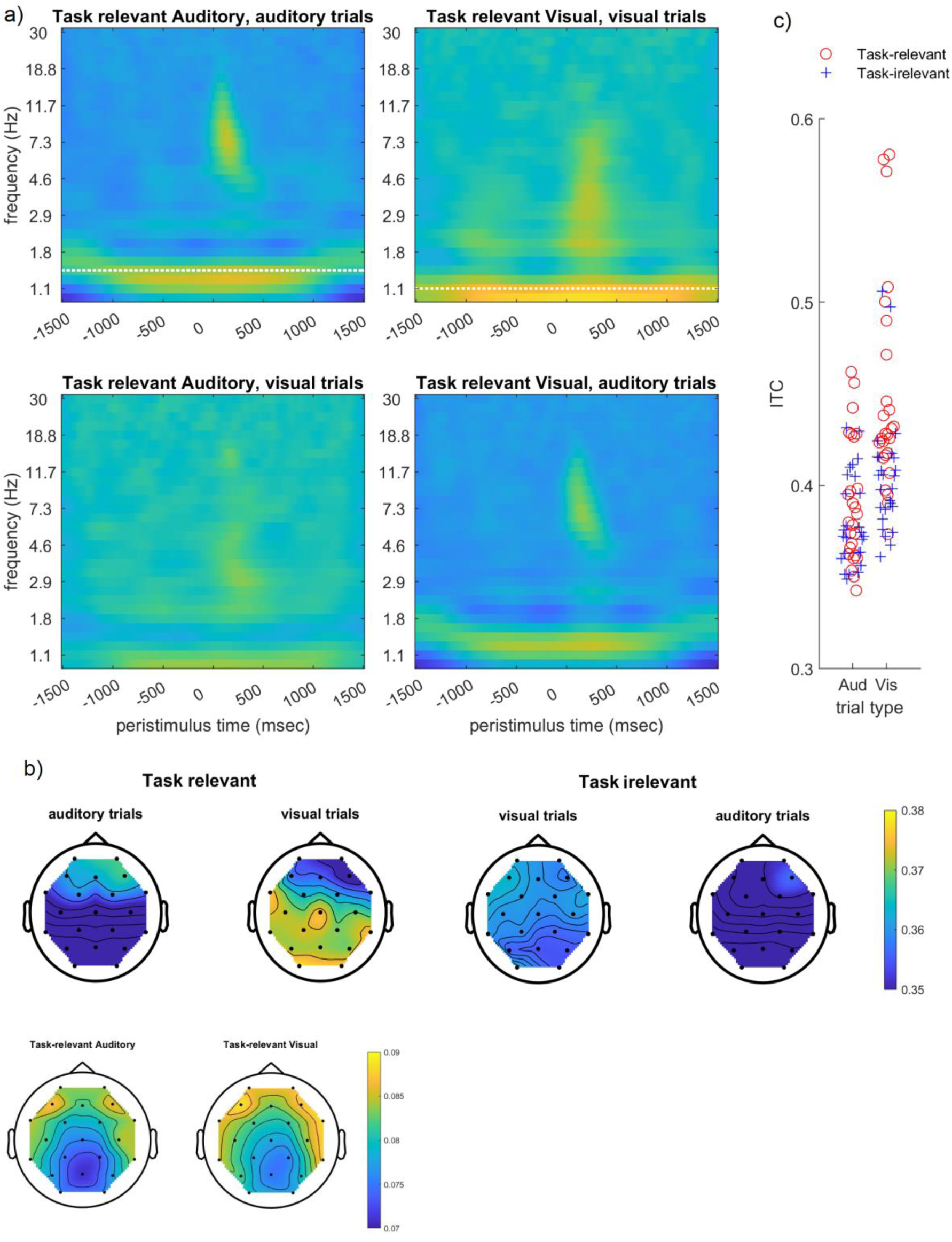
Low-frequency phase entrainment. a) ITC time-frequency spectra, averaged across all electrodes. White horizontal lines indicate the average presentation frequency of auditory (top left) and visual (top right) stimuli. b) Topographies of ITC strength at the peak frequency and the time of stimulus onset. Topoplots for alpha power are displayed in Figure 3-1. c) Average ITC at the driving frequency as a function of task relevance (task-relevant vs. task-irrelevant stimulus sequence) and stimulus type. Figure 3-1. Topoplots of pre-stimulus alpha power distribution for task-relevant stimuli.

### Relation between pupil size, ITC, and prestimulus alpha power

We next inspected the paired associations between pupil size, ITC, and pre-stimulus alpha power. ITC did not show a main effect of pupil bin (**Fig. 4a**; F(1, 992) = 0.16, p = 0.68). However, there was a significant interaction between bin and task relevance (F(1, 992) = 7.74, p = 0.005). Post-hoc linear regression analyses showed that with increasing pupil size the low-frequency oscillations became more entrained to the task-relevant stimulus sequence (β = 0.005, t = 2.22, p = 0.026) while the numerical decrease in entrainment to the task-irrelevant stimulus sequence was nonsignificant (β = -0.003, t = - 1.70, p = 0.089).

**Figure 4.**
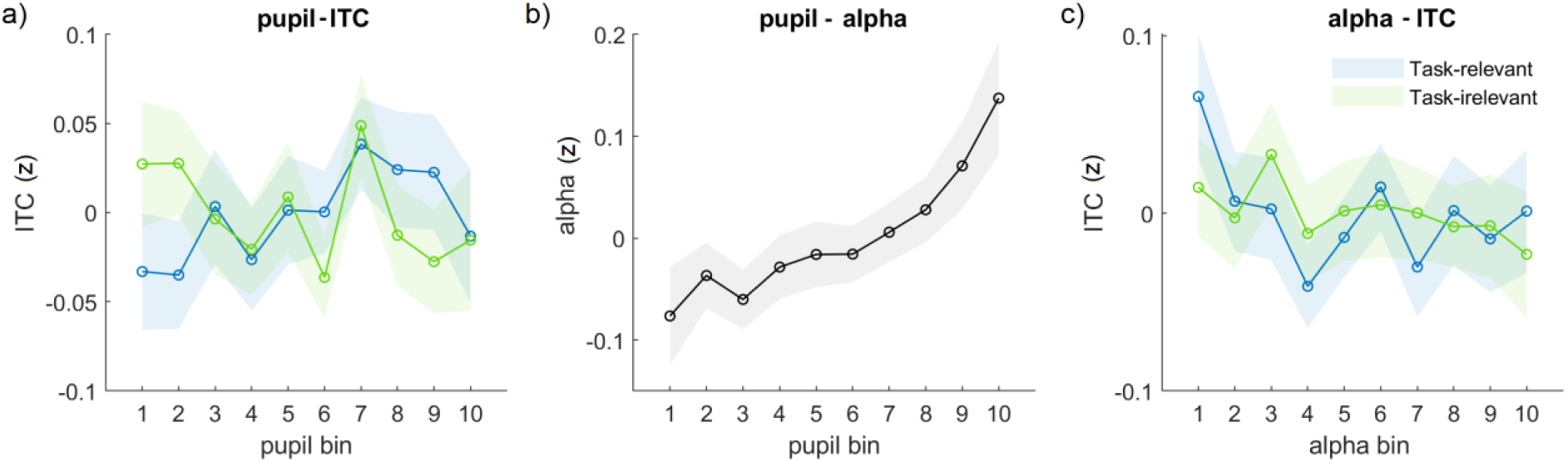
Paired relation between pupil size, ITC and alpha power. Relation between (a) pupil size and ITC; (b) pupil size and prestimulus alpha power; and (c) prestimulus alpha power and ITC. Different colours correspond to the task-relevant and task-irrelevant modality. Note that the distinction between the task-relevant and task-irrelevant modality is only of interest for ITC. There were no significant main effects and interaction effects of stimulus type (visual, auditory, Fig. 4-1) so we averaged across those levels. The significant effects cannot be explained by the number of targets (Fig. 4-2).

**Figure 4-1.**
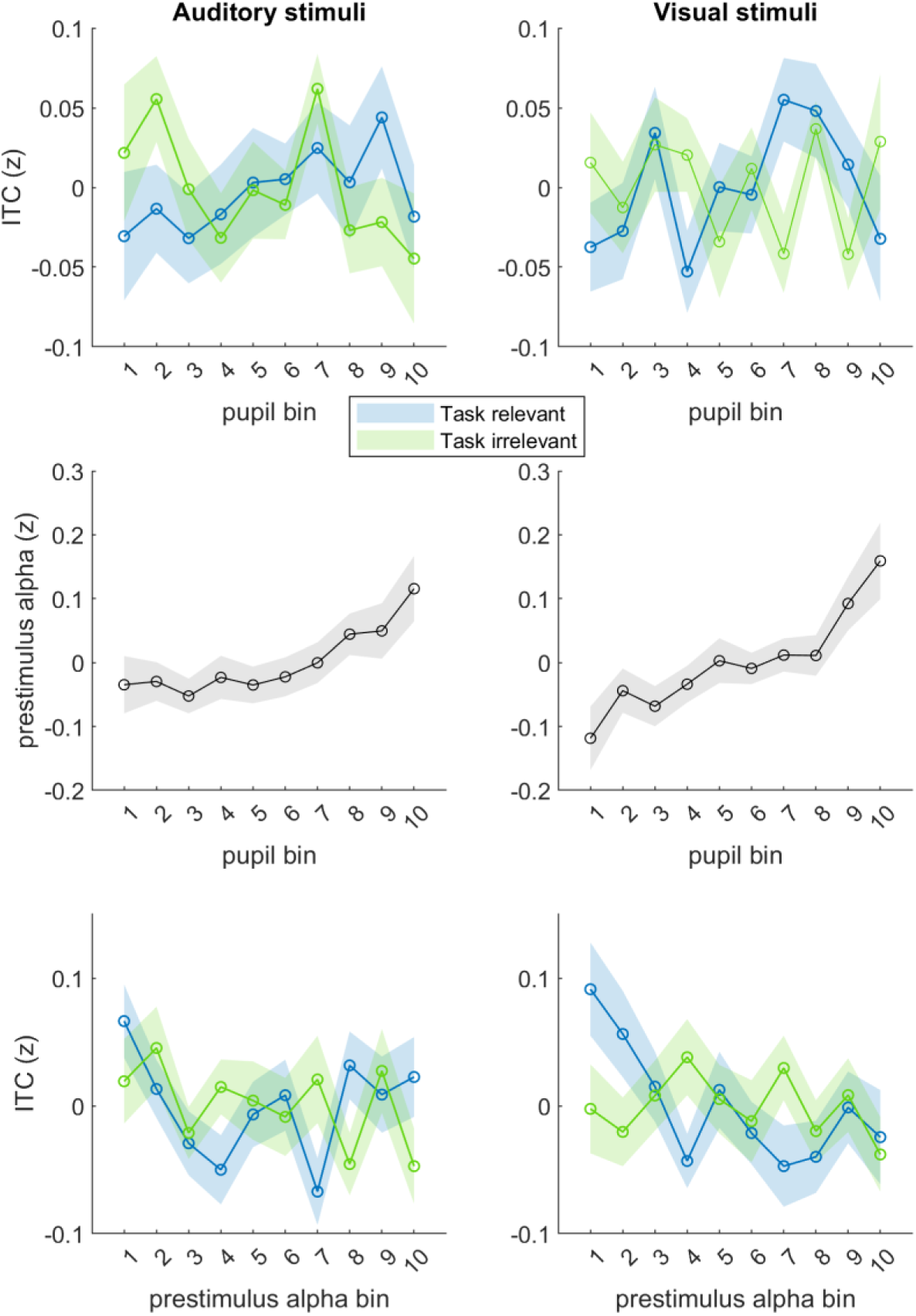
Paired relationships between pupil size, ITC and alpha power separating by stimulus modality. Left column: auditory stimuli in Attend Auditory blocks (i.e., auditory stimuli were task-relevant, blue) and Attend Visual blocks (i.e., auditory stimuli were task-irrelevant, green). Right column: visual stimuli in Attend Visual blocks (i.e., visual stimuli were task-relevant, blue) and Attend Auditory blocks (i.e., visual stimuli were task-irrelevant, green). Top row: pupil size and ITC; Middle row: pupil size and prestimulus alpha power; bottom row: prestimulus alpha power and ITC.

**Figure 4-2.**
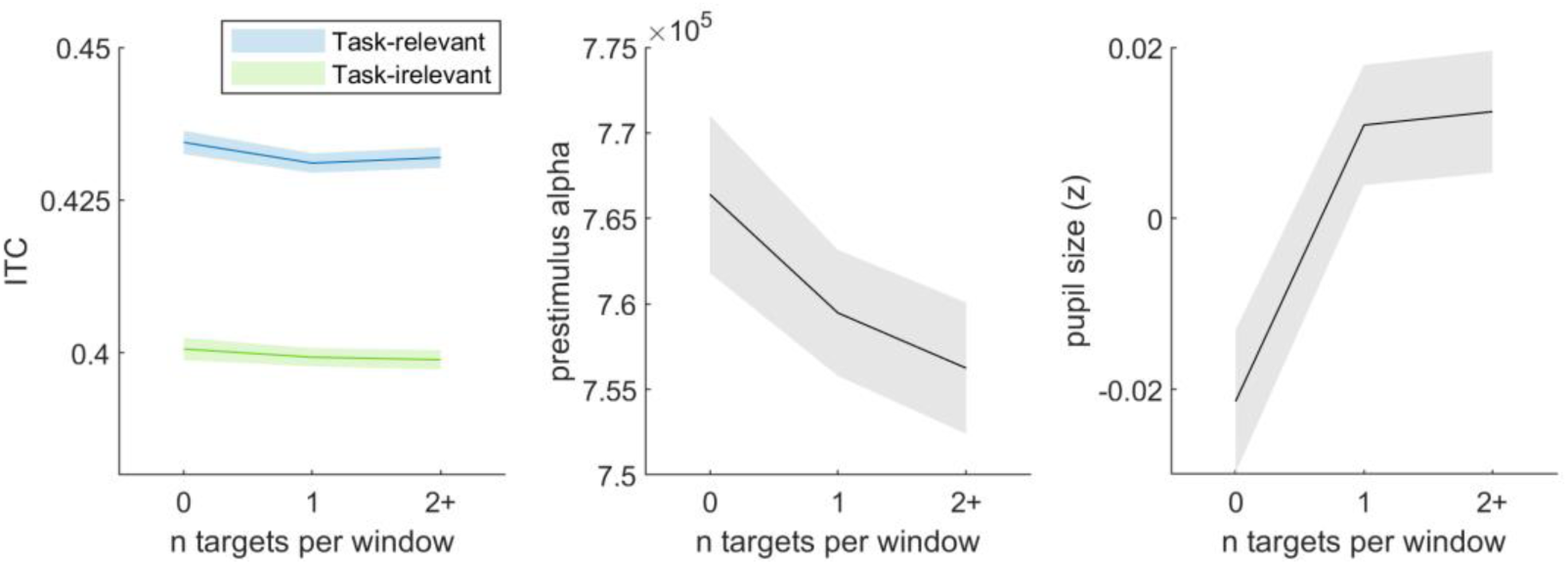
Effect of number of targets on pupil size, ITC and alpha power. We examined if the relationships between our four key dependent variables might be confounded by enlarged evoked EEG or pupil responses to infrequent target stimuli. The temporal overlap between the auditory and visual sequences made it hard to estimate single-trial evoked responses, but we used a more indirect way to address the issue: We assessed if ITC, and average alpha power and pupil size in a given window depended on the number of targets in that window. If two variables were each dependent on the number of targets, then their relationship might be due to this confound rather than slow co-fluctuations in internal state. We found no effect of number of targets (0, 1 or 2+) on ITC in the task-relevant (F = 1.35, p = 0.25) or task-irrelevant modality (F = 0.99, p = 0.41, Fig. 4.2). We found no effect of number of targets on prestimulus alpha (F = 1.47, p = 0.228). We found an effect of number of targets on pupil size (F = 2.91, p = 0.020), with increasing pupil size for increasing numbers of targets. The absence of an effect on ITC and alpha power suggests that number of targets cannot be driving the significant relation between this variable and pupil size in Fig. 4.b.

When looking at prestimulus alpha power, we found a highly significant main effect of pupil bin on alpha power (**Fig. 4b**; F(1, 998) = 108.27, p < 0.001). Prestimulus alpha power directly increased with pupil size, in line with a previous study (Podvalny et al., 2021). Furthermore, ITC showed a significant main effect of alpha power bin (Fig. 4c; F(1, 1032) = 9.00, p = 0.002), with decreasing ITC as alpha power increased (β = -0.004). The interaction with task relevance was not significant (F(1, 1032) = 0.58, p = 0.445).

We did not find any significant main effect of stimulus type (auditory vs. visual) or interaction between stimulus type and pupil/alpha bin (*p* > 0.1; **Fig. 4-1**), suggesting that the key results reported here generalized across perceptual modalities. Also, our key results could not be explained in terms of enlarged evoked EEG or pupil responses to infrequent target stimuli (**Fig. 4-2**).

### Arousal-related modulation of the temporal relation between ITC and alpha power

As an exploratory analysis, we asked whether pupil-linked arousal may modulate the temporal relation between entrainment and alpha power. Specifically, we inspected entrainment and alpha power associated with stimuli of the task-relevant modality in periods centered around pupil maxima and minima (**Fig. 5a**; Montefusco et al., 2022). Although these maxima and minima roughly correspond to the extreme pupil bins in **Fig. 2** and **Fig. 4ab**, they are partly distinct in that they involve moments in which the derivative of the pupil signal, thought to track distinct changes in cortical state (McGinley et al., 2015b; Reimer et al., 2014), changes sign (e.g., from positive to negative around pupil peaks). We found that pupil maxima were preceded by an increase in entrainment, but this effect was transient and small (**Fig. 5b**). More interestingly, a large increase in pre-stimulus alpha was observed prior to pupil maxima, and a large decrease prior to pupil minima (**Fig. 5c**). This indicates that changes in the electrophysiological signatures occurred prior to the changes in pupil-linked arousal. To investigate whether the changes in entrainment and alpha power associated with changes in arousal were linked, we calculated the cross-correlation between these signals around pupil maxima and minima. We found that the temporal relation between entrainment and alpha power was strongly modulated around pupil maxima and minima (**Fig. 5d**): there was a significant positive correlation between ITC and alpha power in time windows around pupil maxima and a significant negative correlation in time windows around pupil minima. Interestingly, while the positive correlation was transient, the negative correlation lasted for a prolonged time (∼18 seconds). Taken together, the results suggest that changes in ITC and alpha power are temporally related, and that this relation is strongly modulated by arousal.

**Figure 5.**
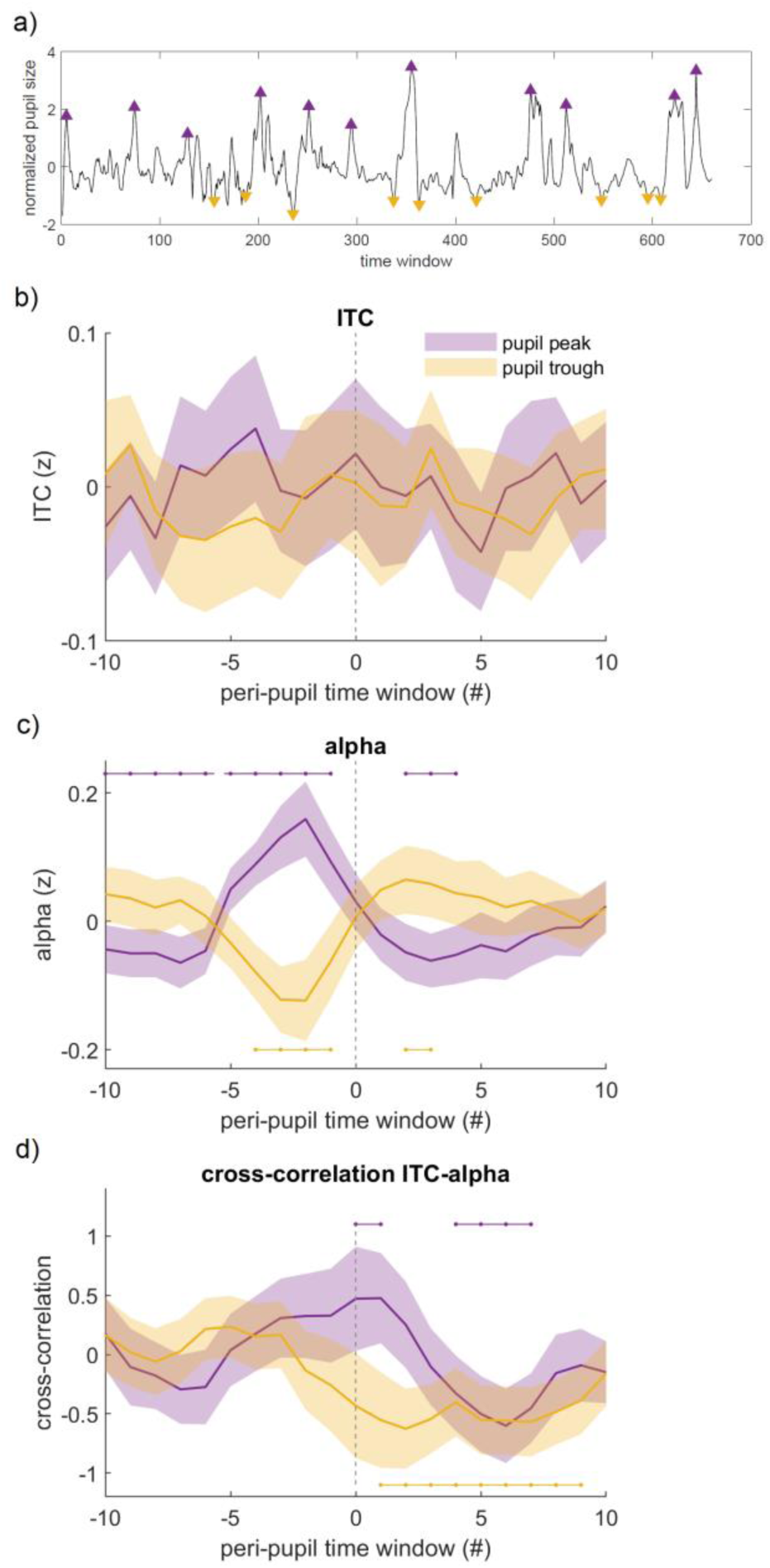
Temporal relation between ITC and pre-stimulus alpha power around pupil maxima and minima. a) Example of one participant’s time course of pupil size and detected maxima and minima marked with yellow and purple triangles, respectively; b) and c) timecourse of ITC and alpha power, respectively, around pupil peaks and troughs; d) cross-correlogram between ITC and alpha power associated with stimuli of the task-relevant modality: around pupil maxima (purple) and around pupil minima (yellow). Negative lags correspond to ITC leading alpha power, while positive lags correspond to alpha power leading ITC. Horizontal lines above and below the time courses indicate significant difference from zero based on permutation testing (p < 0.05).

## Discussion

The goal of the current study was to unravel how variations in the brain’s internal state contribute to fluctuations in neural and behavioral measures of attention at slow timescales. We examined the impact of varying levels of pupil-linked arousal and alpha power on low-frequency phase entrainment. We found that increases in pupil size coincided with stronger entrainment of low-frequency oscillations to the task-relevant stimulus sequence. In contrast, increases in prestimulus alpha power coincided with a *decrease* in oscillatory phase entrainment to both the task-relevant and task-irrelevant stimulus sequences. Furthermore, we found that alpha power and ITC showed a significant correlation, but only in the time windows around pupil maxima and minima, with the sign of the correlation depending strongly on whether pupil-linked arousal was high or low. These results suggest that variability in internal state modulates how the brain responds to external rhythmic stimulation, and that arousal is a key factor coordinating the oscillatory mechanisms of the brain.

### Facilitatory effect of arousal on low-frequency entrainment

A key finding in our study was that increases in pupil size coincided with a selective increase of low-frequency phase entrainment to the task-relevant stimulus sequence. While previous work on arousal-related effects on cortical function has largely focused on processing of single stimuli (either unpredictable or predictable), how arousal may benefit the processing of rhythmic stimulation has been less studied. Previous research has found that diminished phase entrainment is associated with lapses in attention and response accuracy (Lakatos et al., 2016; van den Brink et al., 2014). Our finding suggests that transient decreases in pupil-linked arousal may be one factor underlying these observations. Studies in animals suggest that pupil dilation is associated with changes in neuronal excitability and population activity (McGinley et al., 2015a; Reimer et al., 2014; Vinck et al., 2015), tuning the cortex into an active state (McGinley et al. 2015b). In addition, increased arousal is often associated with better task performance at low-to-intermediate arousal levels, while task performance decreases at very high levels of pupil-linked arousal, giving rise to an inverted-U relation between pupil-linked arousal and performance (Beerendonk et al., 2023; van den Brink et al., 2016; Waschke et al., 2019). Interestingly, **Fig. 4a** shows decreased task-relevant ITC in the time windows with the largest baseline pupil size (bin 10). Our findings suggest that pupil-linked arousal, by shaping the brain’s internal state, affects how the brain processes task-relevant rhythmic stimuli.

Arousal-related neuromodulatory systems have been suggested to play a role in selective attention by setting the level of responsivity (i.e., gain) and excitability of the regions processing single stimuli that are salient or task-relevant (Dahl et al., 2022). In light of our results, one can speculate that while a *phasic* recruitment of neuromodulatory nuclei facilitates the processing of single stimuli, the *tonic* mode may be used for entrainment to sustained stimuli at low frequencies. While future studies are warranted to directly elucidate how neuromodulatory activity affects activity at the neuronal level during rhythmic stimulation, our results pose the arousal-related neuromodulatory systems as a highly flexible tool that can be recruited under different situations.

### Evidence for entrainment versus alpha processing modes

While larger pupil size was associated with a selective increase in low-frequency phase entrainment, larger alpha power was associated with an overall decrease in phase entrainment. These contrasting effects are surprising, given that alpha power and pupil size showed a strong positive relationship, a similar relationship with task performance (i.e., negative quadratic relationship with number of incorrect responses), as well as a clear temporal relationship (**Fig. 5c**): alpha power peaked before pupil maxima and reached a minimum before pupil minima—a similar pattern as in Montefusco et al. (2022). Moreover, both pupil size and alpha desynchronization are thought to reflect activity of the noradrenergic system (Dahl et al., 2022). On the other hand, the partly dissociable effects of pupil size (positively correlated with task-relevant phase entrainment; **Fig. 4a**) and posterior alpha power (negatively correlated with entrainment; **Fig. 4c**) seem consistent with previous work suggesting that spontaneous fluctuations in pupil size and alpha power jointly shape perception but modulate different aspects, i.e. perceptual sensitivity and perceptual bias, respectively (Pilipenko and Samaha, 2024; Waschke et al., 2019). Our study thus offers new evidence on a partial dissociation between these variables as mechanisms modulating attention to rhythmic stimulation.

A recent framework of oscillatory mechanisms proposes the existence of processing modes: an *entrainment mode*, in which sensory brain oscillatory mechanisms are engaged in processing rhythmic stimuli, and an *alpha-dominated mode*, in which the brain decouples from external stimulation (Zoefel and Van Rullen 2017). Our results support this framework in the sense that higher alpha power was related to decreases in entrainment to any (task-relevant and task-irrelevant) stimuli. While evidence for these modes was found in monkeys (Lakatos et al. 2016), here we found this effect also in humans. This is a novel result because it may reflect a general attentional mechanism comprising exogenous (i.e. led by task structure) and endogenous (led by internal state) processes. Furthermore, our results suggest that the shift between these modes is dependent on level of arousal, as we discuss below.

### Arousal-dependent relation between pre-stimulus alpha power and low-frequency entrainment

Our exploratory analyses of the temporal relation between pre-stimulus alpha power, ITC and pupil maxima/minima shed some light on the differential effects of pupil-linked arousal and alpha power: alpha power and task-relevant ITC showed a significant cross-correlation in the time windows around pupil maxima and minima, with the sign of the correlation depending strongly on whether pupil-linked arousal was high or low. We can only speculate why the coupling between entrainment and alpha power differed at the opposite extremes of arousal. One possibility is that the peaking of alpha power and correlation with ITC around pupil maxima reflected activity in brainstem arousal centers, which also drove the peaking of pupil size. Indeed, activity in the locus coeruleus occurs shortly before pupil dilation (Joshi et al., 2016), pupil-linked arousal has generalized effects on the brain that are evident at slower time scales (Shine et al., 2016). On the other hand, the counter-relation around pupil minima may have reflected an inattention mechanism within the alpha band that emerged at low arousal. Indeed, it has been proposed that top-down and primary sensory circuits may contribute to the alpha band with opposing effects (Bollimunta et al., 2008). During low arousal, cortical effects on alpha power may signal basic sensory processes, which are dissociated from pupil-linked arousal effects (Pilipenko & Samaha, 2024). Lakatos and colleagues (2016) reported a slow anti-correlation in nonhuman primates between alpha power and ITC, as discussed above, and a recent study in humans reported a similar anti-correlation with entrainment to auditory stimuli (Kasten et al. 2024). Intracranial measurements in humans also showed a modulation of alpha power by the phase of delta entrainment (Gomez-Ramirez et al., 2011). In terms of mechanisms of selective attention, we speculate that arousal is related to higher entrainment and cognitive contributions to alpha power (Clayton et al., 2018; Sadaghiani and Kleinschmidt, 2016). A slow counter-modulation between sensory alpha power and entrainment, rather than being sustained, may emerge during periods of low arousal. Given that the results in humans and non-human primates suggest the translational value of the findings discussed here, future research should replicate and further disentangle the relation between alpha power and ITC during periods of large and small pupil-linked arousal.

### Limitations and future directions

From a methodological perspective, it is hard to distinguish between true entrainment of ongoing neural oscillations and the consistent phase alignment caused by a sequence of stimulus-evoked potentials (Helfrich et al., 2019; Haegens and Golumbic 2018; van Diepen and Mazaheri, 2018). Nevertheless, mounting evidence shows that phase coherence persists during no-stimulus periods, suggesting that it is at least in part caused by neural entrainment (Bouwer et al., 2023; Zoefel et al., 2018), and intracranial studies in humans support the presence of wide-spread alignment of oscillations to rhythmic stimuli, not restricted to sensory areas (Besle et al., 2011; Gomez-Ramirez et al., 2011).

Previous studies have found within-trial phase-amplitude coupling between the delta phase of entrained oscillatory activity and alpha power over posterior brain regions (Wilson and Foxe, 2020; Wostmann et al., 2016). Here we were interested in fluctuations at slow time scales, similar to Lakatos et al. (2016), because they likely represent variation in the brain’s internal states. To compute the uniformity of phase angles across stimulus sequences we used 8-second sliding time windows. The length of these time windows did not allow for sub-second temporal resolution, limiting our ability to study stimuli-specific relationships between entrainment and alpha power at various levels of arousal. Taken together, our results and previous studies suggest that internally and externally-driven oscillatory mechanisms operate at different time scales.

As our study design was based on Lakatos et al. (2016), we did not strictly counterbalance the order of the Attend Auditory and Attend Visual blocks or the presentation frequencies of the auditory and visual stimulus sequences (i.e. resulting in more auditory trials for the computation of ITC). We could afford this because none of our key hypotheses (i.e., those corresponding to **Fig. 4**) involved a direct comparison between the two perceptual modalities; instead, we recoded the stimulus sequences as ‘task-relevant’ and ‘task-irrelevant’ and focused our analyses on that distinction, as in Lakatos et al. (2016). Although the difference in presentation frequency may have driven the modality effects in ITC, it is interesting that none of the relationships we observed between ITC, alpha power, pupil size and behavior showed systematic differences between the auditory and visual modalities. This supports the assumption that our results reflect fluctuations in internal state rather than modality-specific effects. Future work could assess the relationship between phase entrainment and another internal state variable: the slope of the 1/f shape of the power spectrum –that is, the aperiodic (nonoscillatory) component of EEG activity (Waschke et al., 2021). This slope has been proposed to track the ratio between excitation and inhibition in underlying neural circuitry (Gao et al., 2017), and has been found to co-vary with spontaneous fluctuations in pupil-linked arousal (Pfeffer et al., 2022). In addition, alpha power estimates may be more reliable if aperiodic EEG is taken into account (Cunningham et al., 2023). <u>Conclusions</u>

Neural entrainment is an important instrument of selective attention (Schroeder and Lakatos, 2009). Here we reported delta-phase entrainment to both the visual and the auditory stimulus sequences, with stronger entrainment to the task-relevant modality than to the task-irrelevant modality. This is consistent with previous studies using bimodal stimulation (Besle et al., 2011; Lakatos et al., 2016). Most importantly, we found a significant linear relationship between pupil size and the degree to which entrainment tracked the task-relevant instead of the task-irrelevant stimulus sequence, indicating a facilitating effect of arousal on the perception of rhythmic stimulus sequences. We also found a coupling between alpha power and entrainment around pupil peaks and pupil troughs. Our results give clear indications of a differential modulation of brain oscillatory processes by arousal.

